# Mutation of charged inner pore residues reduce *E. coli* β clamp residency and increase sliding rates on DNA

**DOI:** 10.64898/2026.08.18.745641

**Authors:** Melissa L. Liriano, Micah McCauley, Shreya Ghosh, Dmitry Korzhnev, Thomas E. Wales, Mark C. Williams, Penny J. Beuning

## Abstract

Sliding clamp proteins play central roles in DNA metabolism, including replication and repair. The ring-shaped *E. coli* beta clamp accommodates double-stranded DNA and serves as a platform for proteins involved in multiple DNA transactions. The inner pore of the beta clamp harbors a series of positively charged and polar residues that can bind to the negatively charged backbone of the DNA. These residues are arrayed so that they do not align with the charged phosphates of the DNA backbone. It is hypothesized that this arrangement of these residues provides for the movement of the clamp on DNA as it alternates which residues are bound to the DNA backbone. In this work, we mutated specific charged and polar residues that project into the inner pore of the beta clamp. The beta clamp variants are dimers and have similar thermal stability and in general a similar ability to complement a temperature sensitive strain for growth. One exception was beta-Q149A, which appeared as higher-order species on a native gel although its hydrogen-deuterium exchange pattern measured by mass spectrometry was overall similar to WT beta. These variants all had decreased binding to DNA after loading. Optical tweezers experiments were used to monitor loading on single DNA molecules and measure the rate of beta clamp sliding on DNA. Consistent with the hypothesized role of positively charged residues in the beta inner pore, mutation of one residue resulted in a faster rate of sliding on DNA.

## INTRODUCTION

DNA replication is carried out by DNA polymerases [1, 2]. *E. coli* DNA Polymerase III (Pol III) alpha (α) subunit of the multiprotein replication machinery is the replicative DNA polymerase. Processivity factors, which enable rapid replication, are found in all domains of life and in *E. coli* the processivity factor is the beta (β) clamp, often referred to as the β sliding clamp [3–8]. The bacterial sliding clamp imparts efficiency and processivity on α during genomic replication by encircling DNA and tethering the polymerase so that it does not dissociate from the nascent nucleic acid strand prematurely.

The toroidal β sliding clamp encircles DNA. Structural studies reveal that β is made up of two crescent-shaped protomers that bind in a head-to-tail orientation to form a circular homodimer with two equivalent dimer interfaces (Figure 1) [9, 10]. Each protomer has three distinct domains, with Domains I and III forming each dimer interface. A hydrophobic pocket between Domains II and III serves as the binding site for the clamp’s various binding partners, including all five DNA polymerases in *E. coli*, the δ subunit of the clamp loader complex, mismatch repair proteins MutS and MutL, DNA ligase, and DnaA-related protein Hda [11–18].

**Figure 1.**
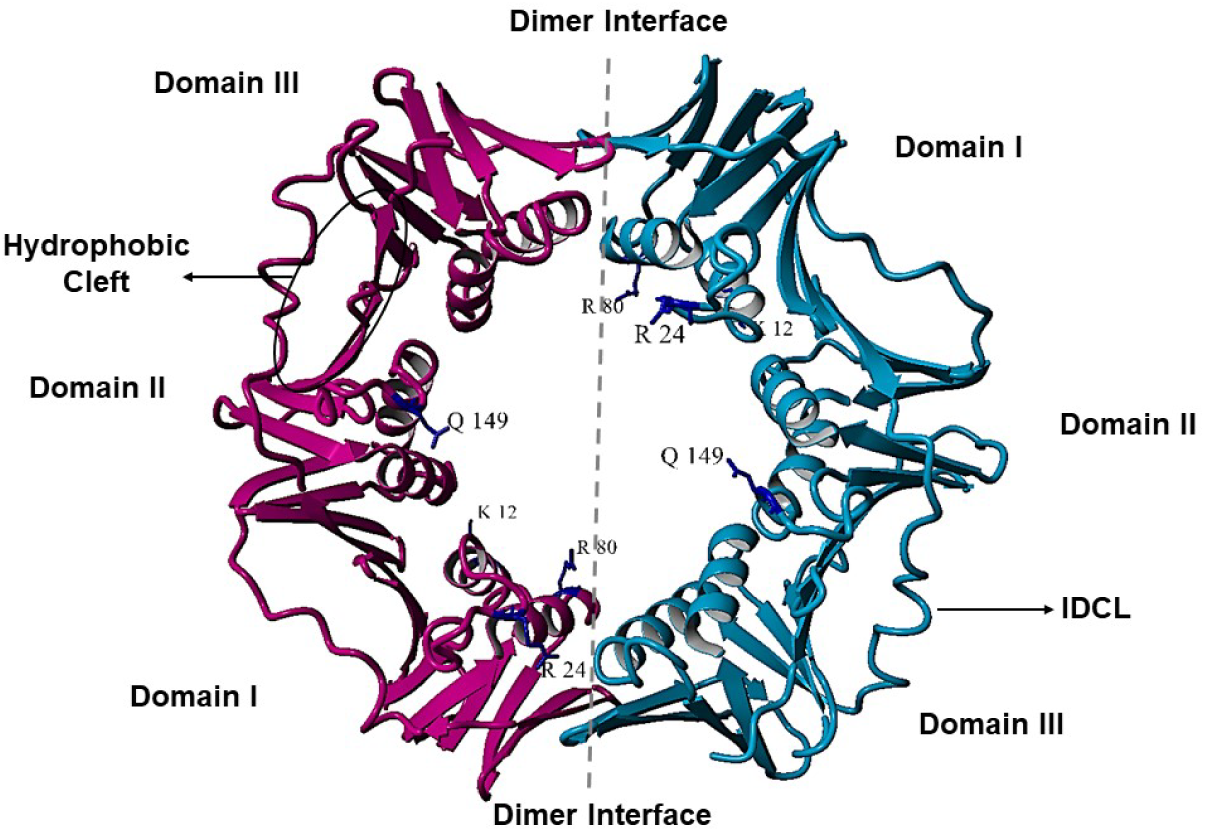
Ribbon representation of β sliding clamp crystal structure (PDB: 1MMI) [30]. Mutation sites are highlighted for residues investigated in this study.

The clamp, especially at the partner protein binding site, is of interest for antibiotic development [19–21]. The inner pore of the β clamp is composed of 12 α helices while the outside of the ring is surrounded by a layer of six continuous beta sheets [10, 22, 23]. Though the overall charge of the β sliding clamp is negative, the α helices lining the central pore of the ring are positively charged [10, 23].

DNA loading of the β sliding clamp is a multi-step process carried out by the clamp loader complex [4, 24, 25]. The clamp loader complex is a member of the AAA+ ATPase family and consists of six proteins: tau (τ), gamma (γ), delta (δ), delta prime (δʹ), psi (ψ), and chi (χ) [6, 25–27]. *In vitro*, the pentameric clamp loader complex consisting of δγ_3_δʹ, also referred to as the gamma (γ) clamp loader complex, can load the β clamp onto dsDNA [10, 28, 29]. In the loading reaction, the γ clamp loader complex first binds ATP, inducing a conformational change that reveals the δ subunit, which binds to β and proceeds to dissociate Domains I and III at one dimer interface, opening the β ring [10, 23, 28–35]. Hydrogen-deuterium exchange mass spectrometry (HDX-MS) analyses show that the β dimer transiently samples open conformations, consistent with δ stabilizing an open clamp [36]. Kinetic studies suggest that spontaneous opening events for the clamp may be too infrequent to facilitate the clamp loader complex capturing an open clamp [37]. Once β binds the γ clamp loader complex, β is preferentially loaded onto a single-stranded(ss)/dsDNA junction with a recessed 3ʹ ends. The loaded clamp encircles DNA and sequential ATP hydrolysis is stimulated in the γ complex, which ultimately dissociates from the clamp and is ejected from DNA [38–40].

When the β clamp loads onto DNA, the DNA contacts several regions of the sliding clamp, including loop I (P20-L27), loop II (H148-Y154), the hydrophobic pocket between Domains I and II, and the inner pore [9, 41]. A co-crystal structure of β on DNA shows that residues R24 and Q149, which are located along the central pore, are among the residues that interact with DNA directly [9]. In the absence of polymerase, a β:DNA co-crystal structure found that the DNA inside the β ring was tilted at a 22° angle [9]. The β residues R24 and Q149 directly interact with the incoming DNA and these residues are not centrally located but instead are found on external loops. It is then hypothesized that R24 and Q149 clamp residues contribute to the tilt observed in the DNA entering the clamp central cavity [41]. The tilt in the DNA may facilitate interactions of β with its myriad protein partners in various biological processes [9, 11–15, 19, 42]. Further analysis of the co-crystal structure provided support for this hypothesis.

First, it was found that the tilted DNA inside the β ring is stabilized by R24 and R197 in one β monomer. Second, the DNA tilt enables positively charged residues R80 and K12, of the second β monomer and on the opposite side of the central pore, to make direct contact with the negatively charged DNA backbone. Since constant breaking and reforming electrostatic interactions at a rate required to tether a polymerase during replication is energetically unfavorable, the sum of these β:tilted DNA interactions may allow β to pause or slow down clamp diffusion, making partner protein/polymerase switching or recruiting feasible [43].

When the β sliding clamp encircles dsDNA, the clamp can diffuse freely along the duplex polymer bidirectionally in a sequence-independent manner [44]. Though central to clamp function, a detailed mechanism of how β slides along DNA is not known. Fluorescence correlation spectroscopy was used to observe the bacterial clamp diffuse on a fluorescently tagged, primed M13 circular DNA. Two types of diffusion, one on ssDNA and another on dsDNA were observed, and the clamp was found to have an average diffusion rate of 10^-14^ m^2^/s, although simulation studies predicted the free diffusion rate over water to be at least 3 orders of magnitude faster [45].

In this study, we examined the influence that the positively charged central pore has on the β:DNA interactions by probing clamp DNA loading efficiency and the rate of clamp sliding on ss/dsDNA. We find that making hydrophobic and negatively charged substitutions to the central pore residues increases the sliding rate and decreases the binding time of clamp variants on DNA relative to wild-type (WT) β.

## MATERIALS AND METHODS

### Protein Expression and Purification

The β clamp protein was expressed from a pET11T vector harboring the *dnaN* gene.

Mutations to the WT β clamp gene K12A, K12E, R24A, R24E, R80A, R80E, and Q149A were introduced using a QuikChange Lightning Site-Directed Mutagenesis Kit (Agilent Technologies). The presence of each mutation was determined through DNA sequencing (Eton Bioscience Inc, Charlestown, MA). For each β variant, protein expression and purification was carried out as described [46, 47].

### Differential Scanning Fluorimetry (DSF)

Melting temperatures (T_m_) of WT β clamp and β variants were determined by incubating samples of 10 μM protein with 10X Sypro Orange dye (Life Technologies) in low-profile, non-skirted, clear 96-well plates at 4 °C, in a final volume of 20 μL, in buffer containing 20 mM HEPES pH 7.5, 50 mM NaCl, 0.1 mM EDTA, 1 mM DTT. Plates were kept at 4 °C and sealed with optical adhesive sheets before placing them in a Bio-Rad C1000 thermal cycler (CFX96 Real-Time System) with fluorescence detected using the channel for fluorescence resonance energy transfer (FRET). Fluorescent signals are reported in relative fluorescence units (RFU). The thermal cycler was set to increase the temperature from 4 °C to 100 °C in 0.5 °C increments every 10 sec. The negative first derivative of melt curves was used to determine protein melting temperature (T_m_). Experiments were performed in triplicate and T_m_ results are reported as the average and error bars indicated the standard deviation.

### Native- and SDS-PAGE

Native gels were pre-run for 20 min at 150 V on ice. Purified β clamp variants (10 µM) were incubated in 1x loading dye buffer (62.5 mM Tris-HCl pH 8.8, 1% Bromophenol Blue, 25% glycerol) and analyzed on 12% Native-PAGE with the resolving gel 320 mM Tris-HCl pH 8.8, 0.1% ammonium persulfate, TEMED. Native gels were run for ∼ 6 h, at 50 V, on ice. For denaturing gels, proteins were incubated in 1x loading buffer (62.5 mM Tris-HCl pH 6.8, 0.01% Bromophenol Blue, 2.5% β-mercaptoethanol (βME), 10% glycerol, 2% SDS), heated for 2 min at 95 °C, and then analyzed by 12% SDS-PAGE. Gels were stained with Coomassie stain, destained in water, and imaged with an iBright FL1000 imager (Invitrogen).

### Hydrogen-Deuterium Exchange Mass Spectrometry (HDX-MS)

Deuterium labelling of the purified proteins (WT β and its Q149A variant) was initiated with a 19-fold dilution into D_2_O labelling buffer (20 mM HEPES, 50 mM NaCl, 0.1 mM EDTA, 1 mM TCEP, pD 7.5 in 99.9% D_2_O). After each labelling time (10 s, 1 min, 10 min, 1 h, 4 h, and 6 h) at 23 °C (room temperature), the labelling reaction was quenched with ice-cold buffer containing 6 M GdnHCl, 150 mM NaH_2_PO_4_, pH 2.1. Undeuterated samples were prepared with the same method as above except using an aqueous equilibration buffer (20 mM HEPES, 50 mM NaCl, 0.1 mM EDTA, 1 mM TCEP, pH 7.5). After quenching, samples were diluted in 0.1% formic acid (1:1) prior to injection into LC-MS followed by digestion online at 15 °C using an in-house pepsin column. The entire HDX chromatographic system was maintained strictly at 0.0 ± 0.1 °C to minimize "back-exchange" (loss of deuterium). Digested peptides were trapped and desalted using a VanGuard Pre-column trap [2.1 mmX 5 mm, ACQUITY UPLC BEH C18, 1.7 µm] for 3 min at 100 µL/min flowrate before being eluted via a 5–35% acetonitrile gradient over 6 min on an ACQUITY UPLC HSS T3 column (1.0 X 50 mm). To ensure data integrity and eliminate peptide carryover between runs, a wash solution of 1.5 M guanidinium chloride, 0.8% formic acid, and 4% acetonitrile was injected through the protease column during each analytical cycle. The final mass spectrometry analysis was performed on a Waters Synapt XS HDMSE mass spectrometer operating in ion mobility mode. The MS runs were calibrated using direct infusion of glu-fibrinopeptide at 200 fmol/µL. Specific instrument parameters include a capillary voltage of 2.5 kV, a sampling cone at 35 V, and a trap collision energy of 4 V. The source and desolvation temperatures are set to 80 °C and 175 °C, respectively. The instrument scans across a range of 50 to 2000 m/z, achieving a high level of precision.

Peptide identification was performed in Protein Lynx Global Server (PLGS) v3.0 using replicates of the undeuterated samples. Peptide masses were identified from searches using non-specific cleavage of a custom database containing the sequence of β clamp protein (Uniprot Id: P0A988). The peptides identified in PLGS software were then imported into DynamX 3.0 (Waters corporation) filtered implementing a minimum product per amino acid cut-off of 0.25, at least two consecutive product ions. The relative amount of deuterium in each peptide for WT β clamp and β Q149A clamp in buffer was determined by the software by subtracting the centroid mass of the undeuterated form of each peptide from the deuterated form, at each timepoint for each condition. The error for determining average deuterium incorporation for each peptide was ± 0.25 Da or less. Relative difference deuterium levels were not corrected for back exchange and thus reported as relative [48].

### Complementation Assay

*E. coli* strain MS120, harboring a temperature sensitive mutation in the *dnaN* gene [49], was used for all complementation experiments. For each variant, MS120 cells were transformed with a plasmid encoding the respective mutated clamp, or with empty vector or a vector encoding WT β serving as negative and positive controls, respectively. Cells were spread onto LB/agar plates containing 100 μg/mL ampicillin. Plates were incubated at 30 °C and 37 °C for 16-18 h. The average colony forming units (CFU) and standard deviations were calculated from experiments performed in triplicate.

### Clamp Loading Assay

Clamp loading reactions were carried out as described using streptavidin (SA) magnetic beads [46, 47]. Briefly, two oligos, one biotinylated 39-mer primer and a 75-mer template, were annealed and conjugated with the SA-beads. The minimal reaction components required for clamp loading were incubated at 37 °C, washed to remove non-specific binding, and DNA loading was assessed at 0 min and up to 60 min of incubation

### Single Molecule C-trap and Flow Cell

For optical tweezers experiments, the β clamp and γ clamp loader complex were labeled with AlexaFluor 633 and AlexaFluor 488, respectively, as described [50]. Experiments were carried out in a LUMICKS C-trap, combining dual optical tweezers and confocal imaging. A quartz laminar flow microchannel cell was used to assemble sliding beta clamps stepwise on a biotinylated DNA hybrid (LUMICKS). Cells were cleaned with a 2% (by volume) solution of Hellmanex III (Helma), or 0.125 M NaOH. A subsequent water rinse was used to verify laminar flow in each channel. The upper (protein containing) channel was then passivated with a 1:20 dilution of 1% casein in PBS buffer. Trapping laser power was reduced to ∼150 mW to minimize solution heating and oxidative damage. Solution buffer consisted of 10 mM HEPES, 100 mM Na+, pH 7.5, unless otherwise noted.

A 1:1000 dilution of 1.0% w/v streptavidin coated polystyrene beads (Spherotech) was flown through the cell until a single bead was caught and verified in each trap. Translating the flow cell brought these beads into a 2:1000 dilution of 20 ng/µL dsDNA (LUMICKS). The 17,853 base pair DNA is labeled with three biotins on each 5ʹ end, a single ATTO 647N label, and two specific nicks. After obtaining and verifying a single tether (via either the fluorescent label or the force-extension curve), the construct was translated into a low salt (10 mM) solution. The DNA segment between the two nicks separates from the tethered strand, and drifts into solution, leaving a hybridized construct of dsDNA and ssDNA. A final translation brought this construct into the passivated channel that contains either 50 or 100 nM of γ clamp loader and β clamp protein in buffer (20 mM HEPES, pH 7.5, 1 mM CHAPS, 7 mM Mg_2_SO_4_ and 1 mM DTT), and 1 mM dATP. During experiments, the tension was held at ∼5 pN to inhibit hairpin formation in the ssDNA.

### Confocal Imaging and Analysis

Confocal images and kymographs were collected in the passivated channel and a weak flow (∼15 nL/s) of protein was maintained during the experiment. Excitation lasers of 488, 532 and 638 nm were used, each with a power of 2-5 µW. Detection is separated by a series of three filters and images are artificially colored as red/green/blue. The blue and red lasers excited the fluorophores attached to the γ clamp loader and β clamp, respectively. The green laser facilitated imaging of the polystyrene beads, which autofluoresce. Pixel resolution was fixed at 40 nm and time resolution for the kymographs was ∼ 50 ms. Confocal images were limited in time by the collection interval, which was typically 1-2 min. For the confocal images, diffraction limited spots corresponded to protein binding. Protein binding in kymographs was revealed as a series of continuous lines. Images were analyzed with Lakeview software (LUMICKS) and further quantified with FIJI. Kymograph tracking software is used to track these binding events [51, 52], and to quantify the binding lifetimes and protein diffusion [53, 54]. The analysis uses the maximum likelihood estimation, which is independent of the bin sizes shown.

## RESULTS

### Inner Pore Mutations Reduce Clamp Stability

The inner pore mutations were chosen to test specific residues that interact with DNA (Figure 1). Melt curves reveal that these point mutations to β clamp polar and charged inner pore residues that directly contact DNA affect thermostability. Melting temperatures of the variants decreased between 1.5-6.5 °C, relative to WT (Figure 2A). All melting temperatures remain well over 37 °C but the decrease in thermostability relative to WT β may indicate slight perturbations in dimer stability.

**Figure 2.**
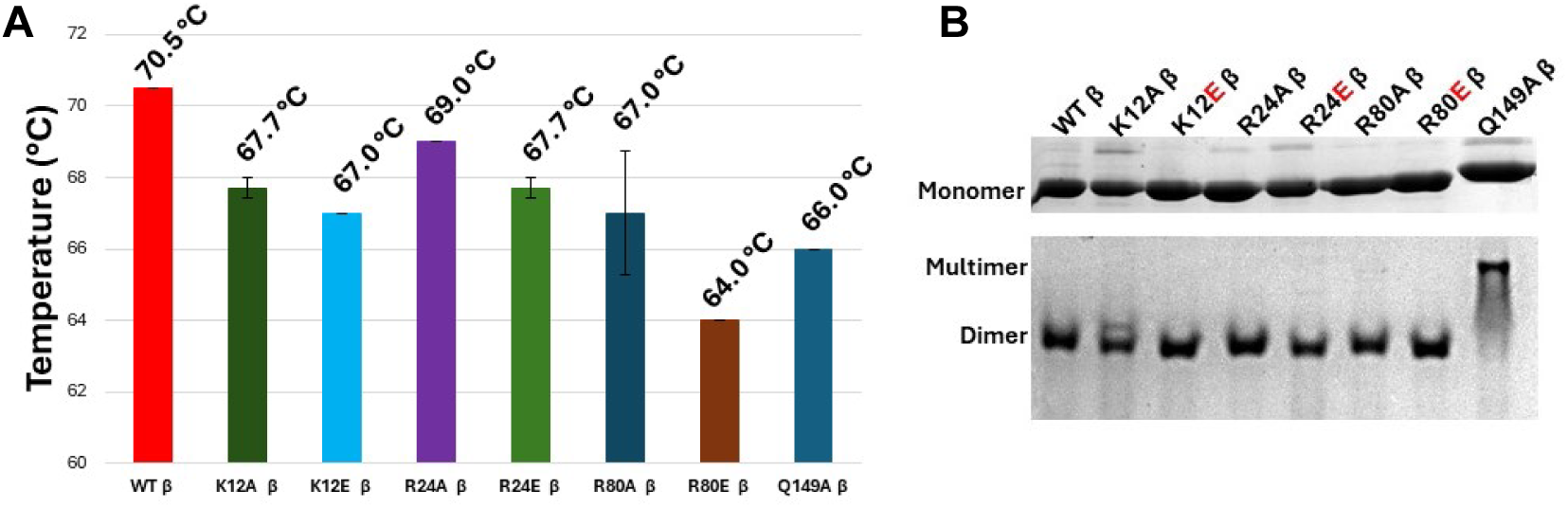
(**A**) Bar graph showing the average melting temperature for each variant. Measurements were performed in triplicate and error bars represent the standard deviations. (**B**) SDS- and Native-PAGE. (top) Denaturing gel showing purified monomers of each clamp variant. (bottom) Non-denaturing gel showing the native oligomeric states of each clamp variant, revealing β Q149A is present as a multimer, [β] = 0.81 mg/mL.

The β clamp variants were analyzed on denaturing and non-denaturing gels. Analysis via non-denaturing gels demonstrates that while most clamp variants maintain their native dimeric state, the Q149A variant exists predominantly as a multimer, suggesting a propensity for aggregation (Figure 2B). HDX-MS analysis of WT β and β Q149A revealed constructs with dynamic, domain-specific architectures (Figure 3A, column 1 and 2, Supplementary Information) with an overall similar global HDX pattern between the two proteins, suggesting the mutation did not introduce gross global changes to backbone dynamics. However, when comparing the HDX-MS profiles of WT β and β Q149A, localized reductions in HDX starting from 10 min after exposure to D_2_O were observed within Domain II and the C-terminal end of Domain III, spanning residues 245–262 and 359–366, respectively (Figure 3A). These observations support a notion that the Q149A mutation induces only a localized conformational change that is marked by the reduced dynamics of selective regions of the backbone, including the C-terminal end of Domain III (Figure 3B), potentially biasing dynamics towards a non-functional oligomerization pose.

**Figure 3.**
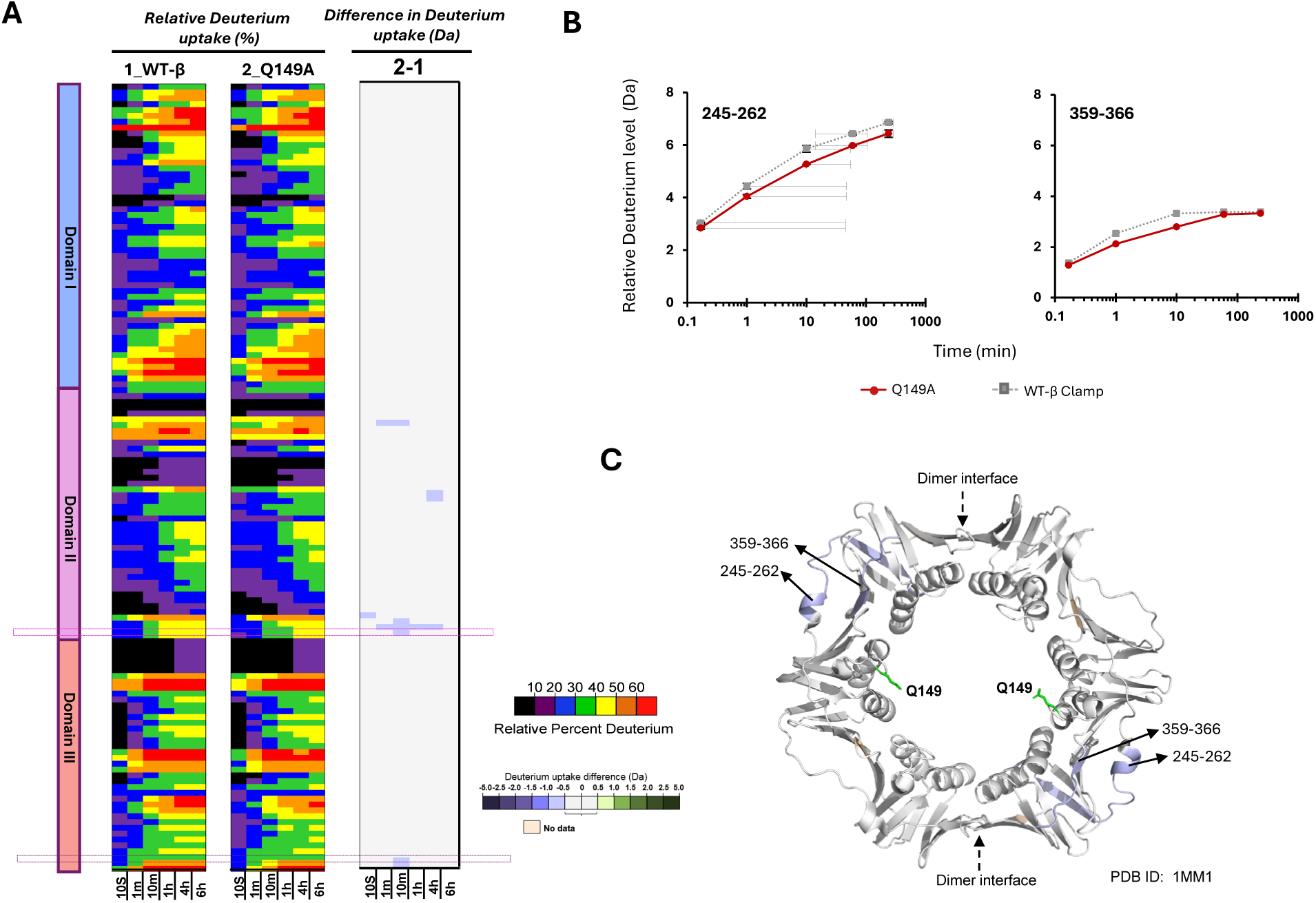
(**A**) Relative deuterium uptake for WT β and β Q149, with the scale shown bottom right, followed by the difference in deuterium uptake. (**B**) Relative deuterium uptake levels for the peptides highlighted in A are plotted as a function of time (min). (**C**) HDX data mapped onto the β structure (PDB: 1MM1) [30], relative to the location of Q149. Peptide coverage map is Figure S1 and deuterium uptake values are in Table S1.

### Most Variants with Mutations to Inner Pore Residues Confer Growth at Non-permissive Temperature

To assess if the β variants are functional clamps, the *E. coli* MS120 strain was used to carry out complementation assays. The MS120 cell line harbors a defective gene for the β clamp (*dnaN159*), which confers a temperature sensitive phenotype for growth at temperatures above 34 °C [49, 55]. At 37 °C, colony formation was observed for all variants tested. Variants with inner pore mutations, while functional in this assay, did not confer growth to the same degree as WT β (Figure 4).

**Figure 4.**
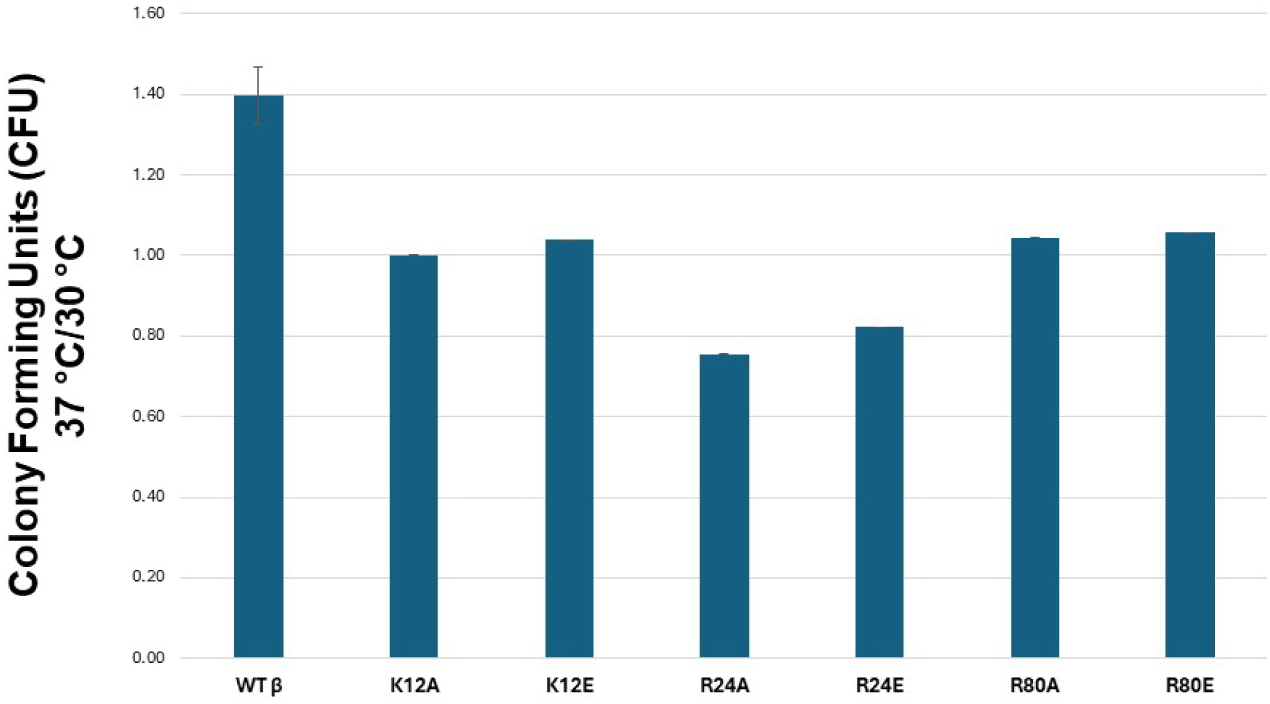
Complementation Assay. Colony forming units (CFU) were calculated as a ratio of the number of colony-forming units at 37 °C vs 30 °C. The β Q149A was not included here given its perturbed dimer formation (Figure 2).

### Mutations to Inner Pore Residues that Interact with DNA Result in Altered β Loading

Commercial streptavidin (SA)-magnetic beads were used to perform DNA loading assays (Figure 5A). Immunoblots with an anti-β primary antibody were used to detect the amount of β present at different steps in the reaction. Band intensities at P0 correspond to the amount of β loaded on DNA after three washes (W), showing that all β variants load onto DNA (Figure 5A). To confirm that clamp variants interact with the oligos adhered to the beads, DNA loading experiments were repeated in the absence of DNA and showed that β Q149A interacts with the SA beads (Figure 5B).

**Figure 5.**
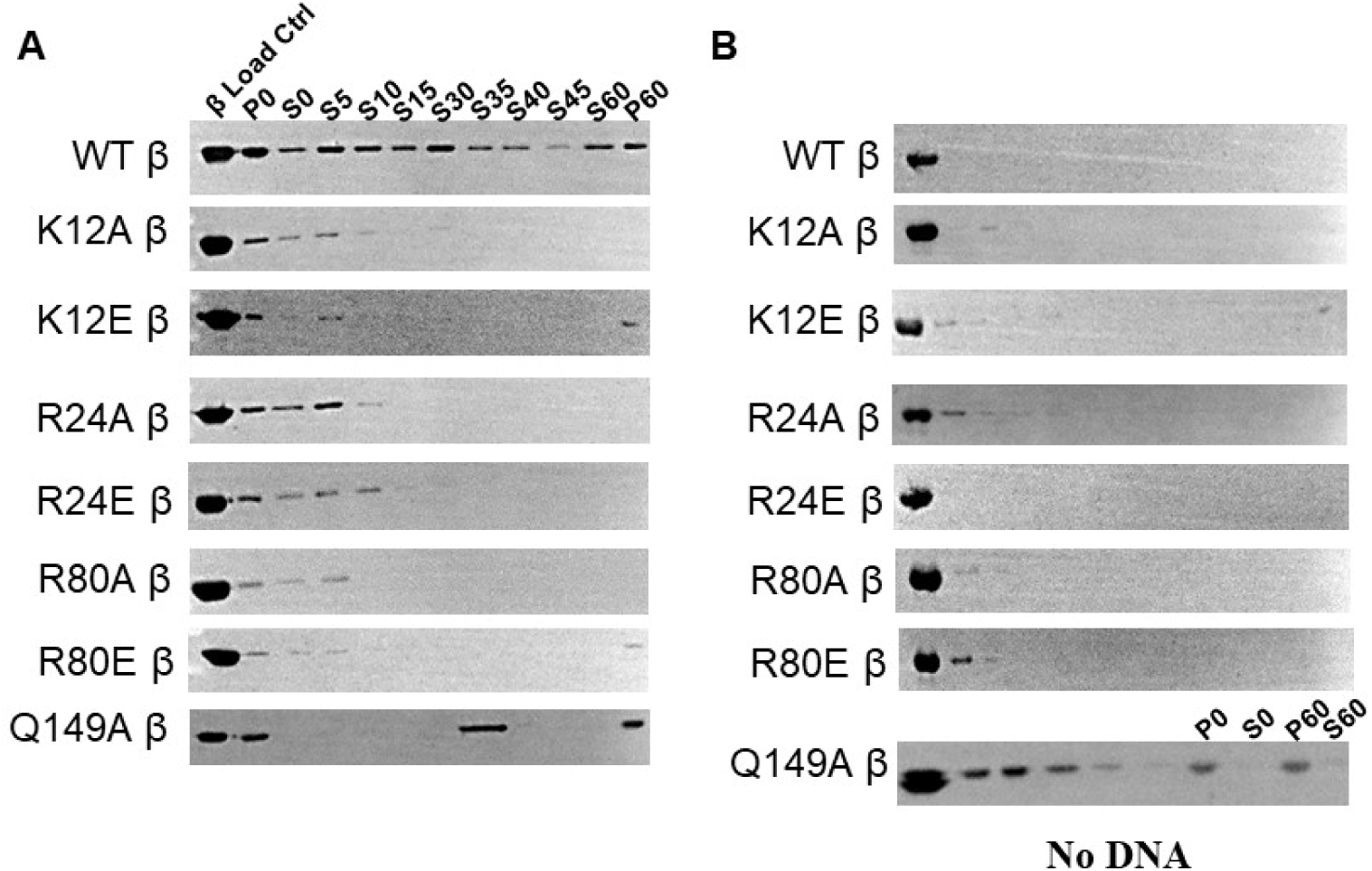
Immunoblots for DNA Loading Assays of β Variants. (**A**) DNA loading of clamp variants. Clamps are detected by an anti-β primary antibody. “P0” corresponds to the amount of clamp loaded onto DNA after washes. β bands in “S” lanes corresponds to the amount of clamp in the supernatant that has unloaded at indicated timepoints (min). (**B**) As a negative control, loading assays were carried out in the absence of DNA, showing that the β Q149A variant and to a lesser extent, the β K12E variant, bind the commercial streptavidin-beads.

Differences in β band intensities in the P0 lanes indicate that the β variants with inner pore mutations, except β Q149A, load onto DNA with different efficiencies. Furthermore, the β variants also appear to have shorter lifetimes on DNA relative to WT β, as revealed by the amount of β detected in the supernatant “S” lanes at different timepoints, with most of the β variants with an inner pore mutation unloading from DNA in less than 30 min. In contrast, WT β remains bound to DNA after 60 min of incubation.

### Alpha (α) Polymerase Enhances Clamp:DNA Interaction

The bacteriophageT4 clamp, gp45, is a homotrimer with significantly weaker trimer interfaces, requiring interaction with its cognate polymerase to remain associated with DNA [56]. To further probe the reduced loading and lifetime on DNA of the β variants observed here, we carried out DNA loading assays in the presence of the Pol III α subunit to examine whether β variant:DNA interactions would be stabilized. In the presence of the α polymerase, β variants show less unloading between 5-60 min (S5-S60, Figure 6A), compared to the amount in the absence of polymerase (Figure 5A). Also, the lifetime of β on DNA is extended in the presence of α polymerase, with β variants K12A, R24E, and R80E having detectable amounts of β present in the pellet and therefore still loaded on DNA at 60 min of incubation. These experiments were repeated in the absence of DNA to confirm that the clamp loading detected was truly an interaction between the β variants and the primed DNA conjugated to the SA beads, which was the case (Figure 6B). In the absence of DNA, a band is detected in the P60 lane for β K12E, indicating an interaction of this variant with the SA-magnetic beads.

**Figure 6.**
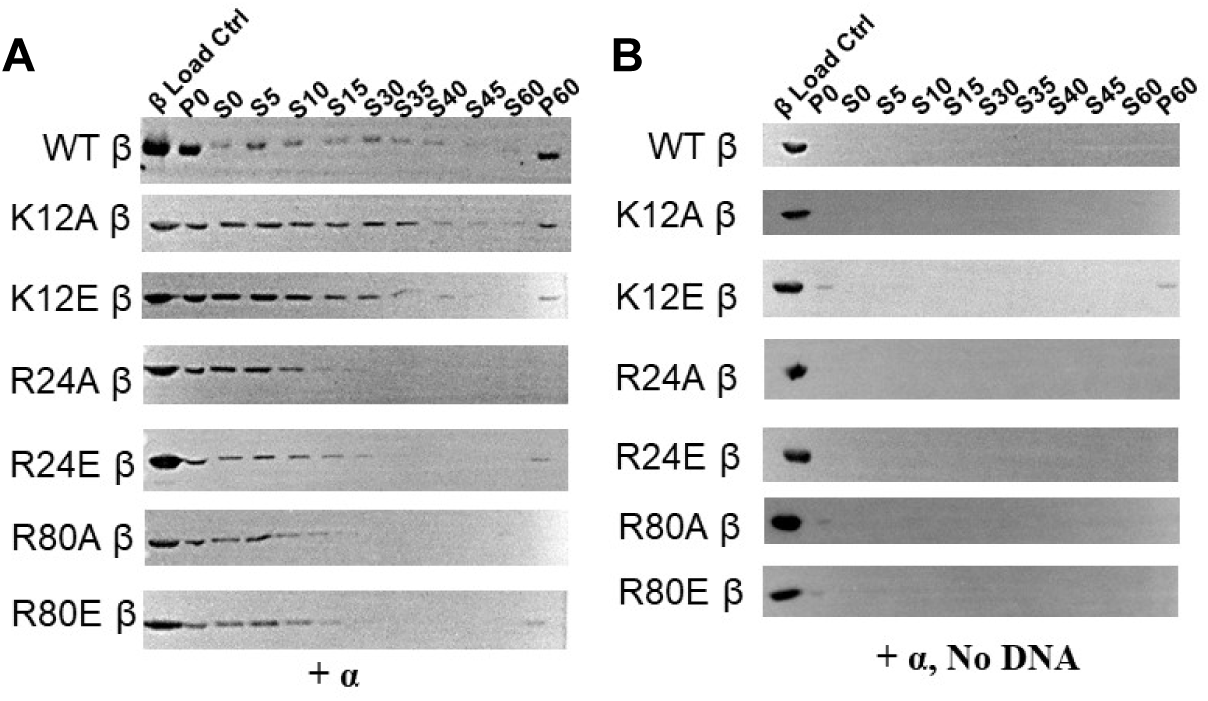
**(A)** Immunoblot of DNA loading assays with all reaction components in the presence of alpha polymerase. **(B)** Immunoblot of DNA loading assay in the presence of alpha polymerase, without DNA [negative control for (B)].

### Visualizing Clamp Loading and Sliding on DNA

To further probe β interactions with DNA, optical tweezers with fluorescence detection were used. Nicked constructs were extended with force in low salt (Figure 7). At an applied tension of >60 pN, dsDNA overstretches and unstacks, and base pairing is lost. The length between each base increases from ∼0.34 nm to ∼0.58 nm [57, 58]. Extending the construct to 9.0 microns in length means that not all of the DNA is overstretched to prevent the unmasking of induced nicks common in handing of long DNA. However, the segments between the two nicks is A-T rich, compared to the flanking sequences that are G-C rich. Furthermore, at only 10 mM Na^+^, significant strand repulsion facilitates the release of the strand between the nicks. As the tension is relaxed, the overall contour length of the hybridized ss/dsDNA construct is ∼7 µm, compared to the ∼6 µm length for fully dsDNA (Figure 7). Thus, the change in length is a reliable marker. The 3ʹ ss/dsDNA junction is marked by a single fluorescent dye, though it was not always visible, especially among the background of the protein solution.

**Figure 7.**
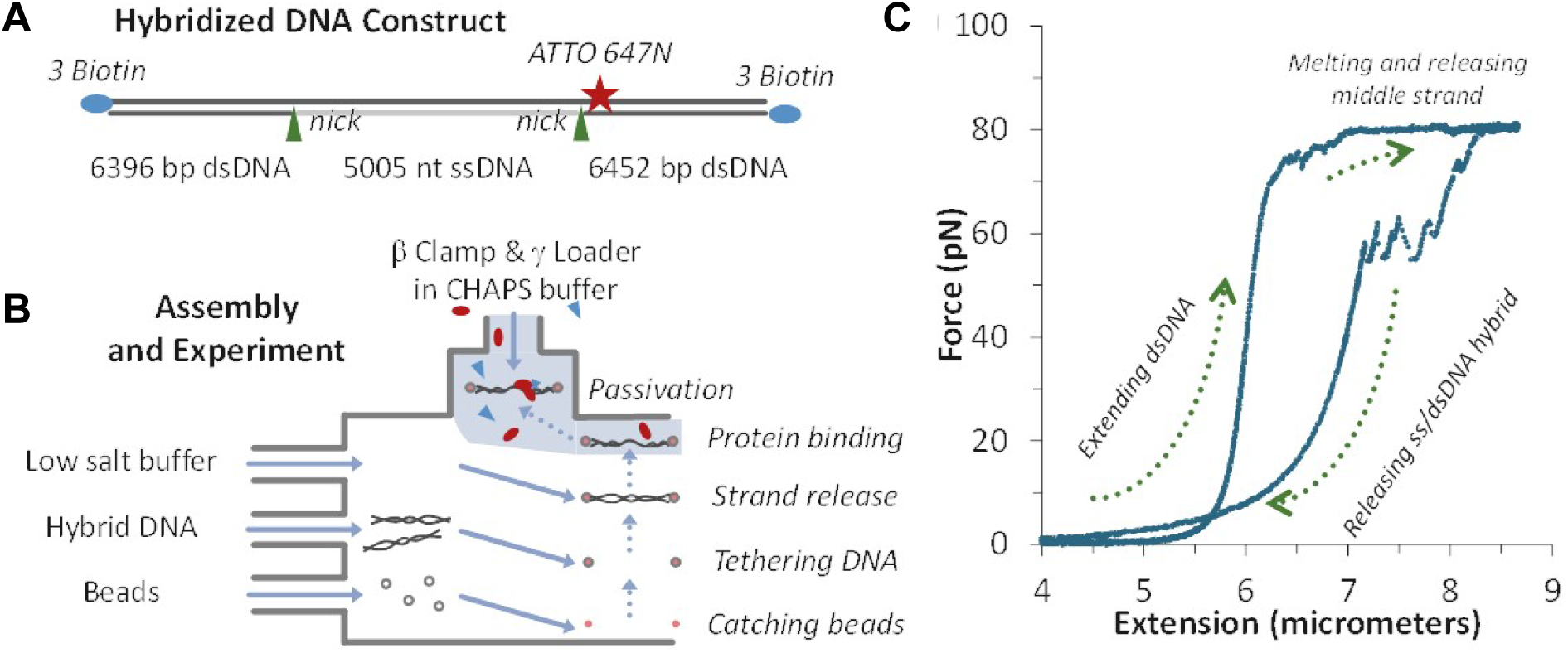
Assembling the beta clamp on hybridized DNA. **(A)** Labeled hybrid DNA is nicked at spots shown. Once caught between two beads and force extended, the middle strand (gray) is released under tension. **(B)** Experimental assembly in a laminar flow cell begins when a hybrid construct is tethered between two trapped beads, and force extended in low salt to create the hybridized strand which is then translated into a solution of labeled clamps and loaders. **(C)** Force extension data of DNA construct, which relaxes to a new contour length corresponding to a hybridized ssDNA/dsDNA.

A series of confocal images initially reveal signals from both the γ clamp loader complex and the WT β clamp, with fluorescent emission centered around 520 nm and 670, respectively (Figure 8). These diffraction-limited images may be seen distinctly. The location of the binding site appears to be consistent with the location of the 3ʹ ss/dsDNA junction. After several min, the signal from the loader disappears, indicating unbinding. At this time, slow movement of the β clamp is evident. The clamp remains on DNA for an average of 8 min before dissociating from DNA, which is consistent with the previously reported 9-min clamp lifetime on DNA in a single-molecule FRET study that captured clamp diffusion [59]. Eventually, the clamp dissociates from the DNA construct, as evidenced by the loss of fluorescent signal on the DNA, though some degree of photobleaching may also occur. To overcome the time resolution limit inherent in confocal image collection, kymographs were also imaged, with 50-ms resolution (Figure 8E).

**Figure 8.**
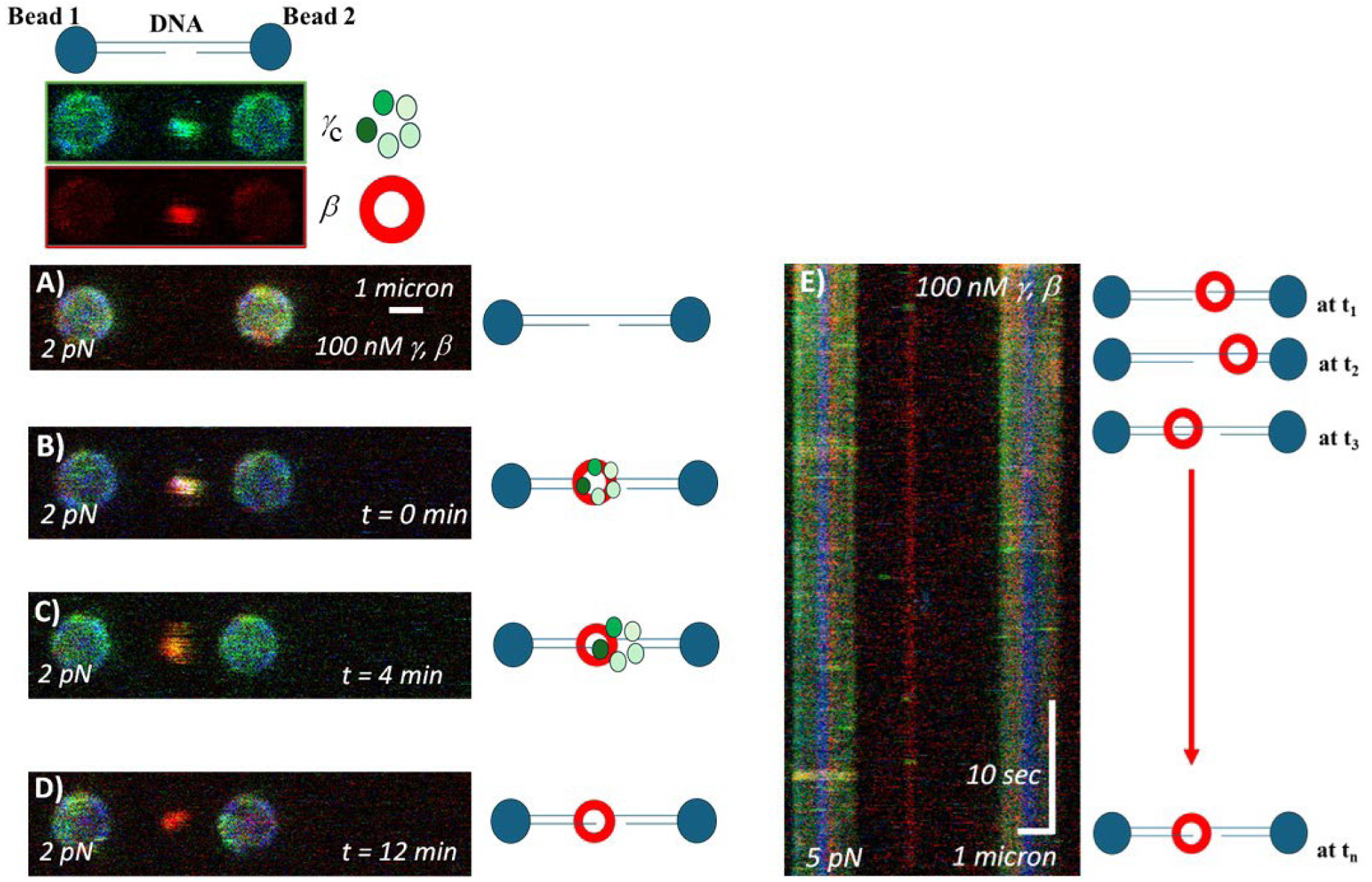
Visualizing loading and sliding. **A)** Two beads, with a hybridized template (not visible) tethered between them. **B)** Protein binding evident from fluorescence signal at the ssDNA/dsDNA junction. Separating the discrete emission channels confirms the distinct presence of γ clamp loader complex (green) and WT β clamp (red). **C), D)** The γ complex dissociates and β moves along the template. **E)** Kymograph of β clamp movement and eventual loss. Increasing time is shown from top to bottom of the image. Results are summarized in Table 1.

**Figure 9.**
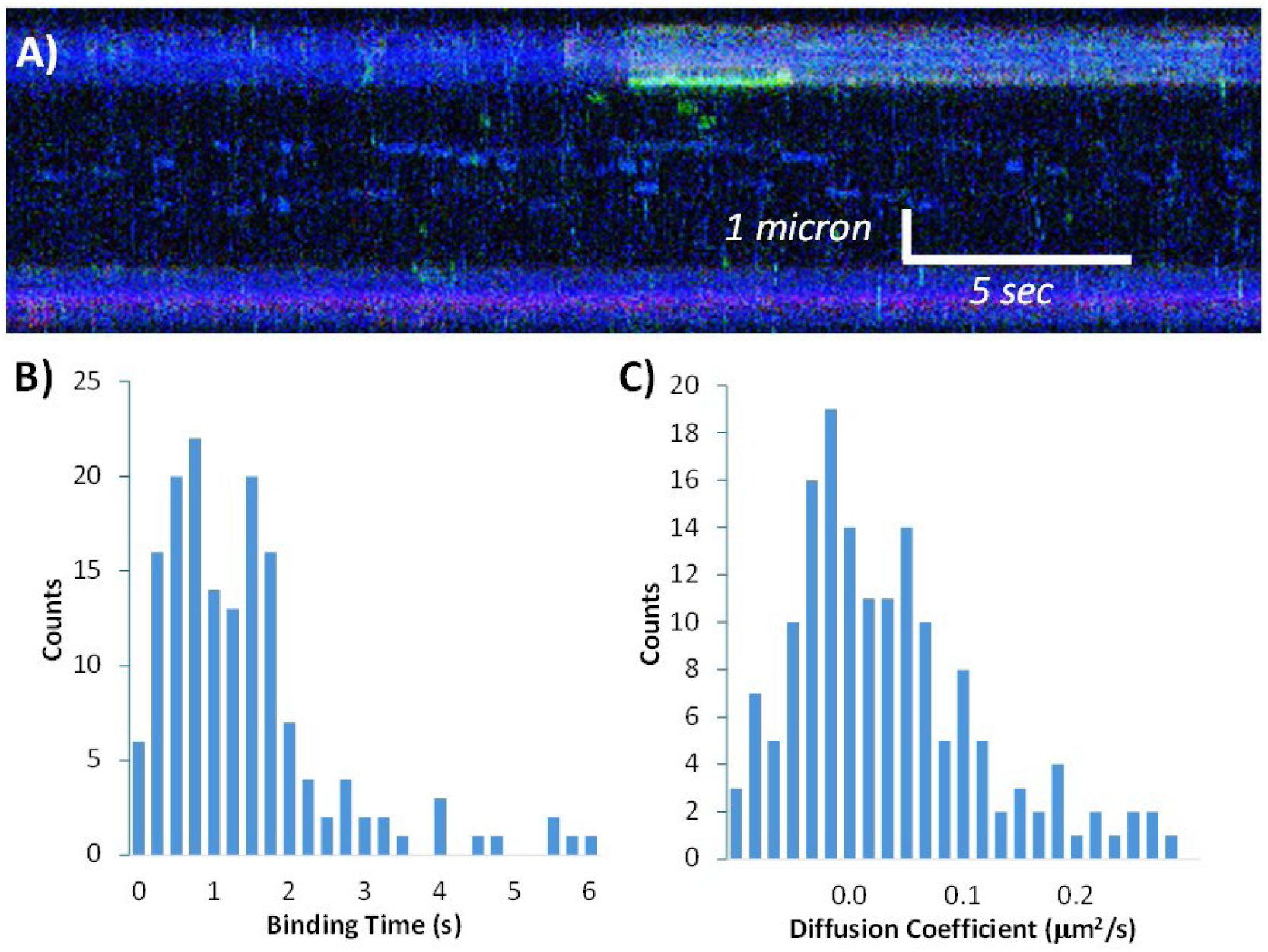
Characterizing γ loader interactions with ssDNA. **(A)** Kymograph of transient binding of γ loader to ssDNA. **(B)** Histogram of binding time, giving a lifetime of 1.26 ± 0.07 s (and a corresponding dissociation rate constant of 0.79 ± 0.04 per s). **(C)** Histogram of measured diffusion coefficient, with an average value of 0.027 ± 0.004 μm^2^/s.

**Table 1.**
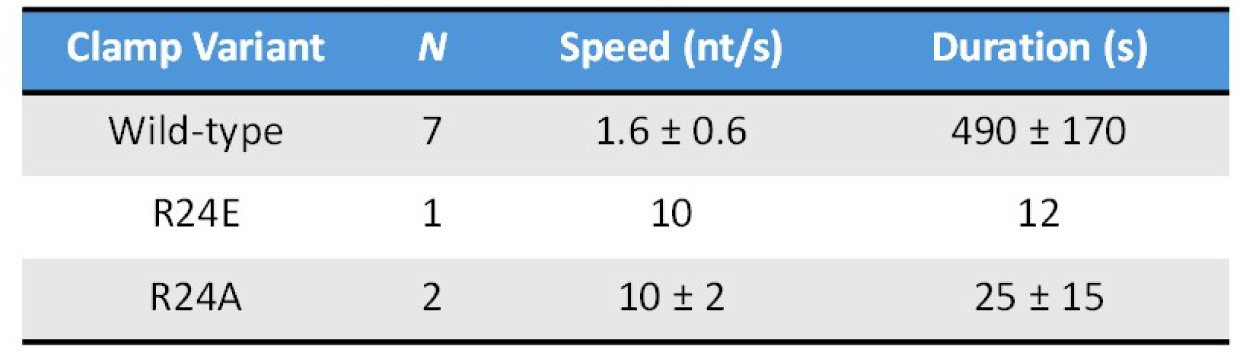
Summary of beta clamp movement on DNA Average measured speed and binding time of beta clamps on DNA as described in the text, with errors due to multiple measurements where possible.

| Clamp Variant | *N* | Speed (nt/s) | Duration (s) |
| --- | --- | --- | --- |
| Wild-type | 7 | $1.6 \pm 0.6$ | $490 \pm 170$ |
| R24E | 1 | 10 | 12 |
| R24A | 2 | $10 \pm 2$ | $25 \pm 15$ |

Roughly half of the clamps move along the gap that corresponds to the single stranded DNA, while half move along the double stranded region. In previous single-molecule FRET clamp diffusion studies, the β clamp was observed also to diffuse on both ss and dsDNA [43]. In that work, single-stranded DNA binding protein (SSB) was included in clamp diffusion experiments to restrict clamp movement along dsDNA; the clamp was observed immobile at 3ʹ end junctions in the presence of SSB [43]. Finally, though γ loader complex and β clamp release are observed in Figure 8, this was not observed in every replicate, possibly due to time resolution limitations or photobleaching. Both the rate of movement and the overall duration of the clamp binding were measured (Table 1).

The site directed mutations made to clamp residues that interface with DNA were also tested in these experiments. Clamp variants β R24A and β R24E have comparable diffusion rates and both move faster along ds and ssDNA than WT β (Table 1). In addition, the overall β:DNA binding time for these variants is significantly reduced relative to WT β (Table 1). This is consistent with the shorter time on DNA observed in DNA loading assays for β R24A and β R24E, relative to WT β (Figure 5A).

### Dynamics of Gamma (γ) Clamp Loader Complex on ssDNA

The γ clamp loader complex was observed to bind to ssDNA (Figure 8). Binding of γ complex to DNA is brief, with a lifetime of 1.26 ± 0.07 s, which corresponds to a dissociation rate constant of 0.79 ± 0.04 per s, and diffusion along ssDNA was also slow, with an average value of 0.027 ± 0.004 μm^2^/s. The γ complex binding to dsDNA was not observed.

## DISCUSSION

### Removing Positive Charge from Clamp Regions that Interface with DNA Leads to Impaired DNA Loading and Binding

The standard model of clamp loading provides a description of the active process by which sliding clamps are assembled onto DNA [4, 25, 28, 29, 31–35, 60, 61]. Recent cryo-electron microscopy (cryo-EM) studies have refined this view, showing that the loader complex undergoes distinct conformational shifts: initially expanding to open the clamp followed by its constriction to facilitate its closure around the double helix [62, 63]. Crucial to this mechanism are specific residues within the β inner pore, such as R24, R80, Q149, Y153, and Y154, which contact the DNA template to ensure efficient loading [4, 9, 12, 25, 41, 43, 60]. To assess the importance of these residues to β:DNA interactions, we introduced mutations to alanine or glutamic acid at K12, R24, R80, and Q149. The expectation is that the Ala mutations would compromising loading activities due to loss of ionic interactions between β and DNA. Likewise, a Glu mutation to residues that directly contact DNA would introduce repulsion of the negatively charged DNA backbone. For the most part, our results were consistent with expectations as R24A/E, K12A/E, and R80A/E variants resulted in reduced DNA loading efficiencies and reduced lifetime once loaded, relative to WT β.

Related to β Q149A, previously published DNA loading assays have shown that a β clamp variant with a five-residue Ala substitution (H148A-R152A), β^148-152^, has impaired loading and poor retention on DNA, however when loaded, the variant is proficient for supporting replication [64]. Intriguingly, a crystal structure of the five-residue Ala substitution (β H148A-R152A) shows that this variant is an intact dimer similar to WT β, with an RMS difference of 0.71 Å [64]. However, in one protomer, the positions of Y153 and Y154 are shifted dramatically relative to their positions in the other protomer or in WT β [64], consistent with changes observed in HDX-MS here and with previous mutation-induced asymmetry observed in β [46, 47, 65]. Another study investigating DNA loading efficiencies with bead-based assays of singly mutated β Q149A or β R24A Q149A showed the variants to be defective in loading onto DNA [9]. Furthermore, elongation speed supported by β Q149A during primer extension was determined to be similar to WT [9]. In our study, using a similar bead-based assay to assess DNA loading, we found the β Q149A variant to have anomalous loading activity with a direct interaction with SA-beads, as observed with other β variants [47]. HDX-MS revealed that β Q149A shows localized protection from HDX in Domain II compared to WT β, providing evidence of modified Domain II dynamics and suggesting that its perturbed state presents a surface mimicking known SA-binding motifs [66]. The Q149A mutation likely disrupts essential inner-pore DNA contacts while promoting a misfolded or dynamically altered architecture that favors sequestration into a multimeric state, as observed by non-denaturing gel electrophoresis. These structural and backbone dynamics shifts prevent the loader complex from executing the required conformational expansion but also likely occludes the critical protein-interaction surfaces in Domain III, rendering the clamp biologically inert.

In the presence of polymerase α, loading efficiencies for the β variants improved and retention on DNA increased. In the presence of Pol III polymerase α, a β:DNA co-crystal structure revealed that the DNA is perpendicular to the β clamp [67]. In this orientation, there is a significant distance between the β amino acids and the dsDNA backbone, not allowing for direct electrostatic interactions and minimizing repulsions. In another crystal structure over 600 ordered water molecules were found hydrogen-bonded to basic and hydrophilic residues lining the inner pore region of the β ring, suggestive of a water-mediated interaction between the β clamp and a nascent dsDNA substrate [30, 43]. With water as a mediator, it is then possible to accommodate the negatively charged regions of the central pore near the DNA backbone [43].

### Mutations to β Inner Pore Residue R24 Leads to Increased Diffusion Rate but Shorter Residency on DNA Relative to WT

During *E. coli* DNA replication, the main polymerase can elongate daughter strands at a speed of 500-1000 nt/s when associated with the β sliding clamp [68]. The sliding clamp is free to move along DNA as it tethers the Pol III polymerase from behind as α adds nucleotides.

Simulations of β predict a free, frictionless diffusion rate of 5.7x10^-11^ m^2^/s for the β clamp [45]. Experimentally, the diffusion rate was found to be significantly slower with D = 10^-14^ m^2^/s, or ∼ 40 nt/s [45].

In our optical tweezer study, we determined the WT β clamp moves along DNA with a speed of 1.6 nt/s ± 0.6, which is significantly slower than the experimental rate previously reported. The reduced drift indicates a large amount of friction is experienced by WT β clamp on DNA. The source of this friction can be strong ionic interactions with the DNA backbone, water viscosity in the medium in which loading reactions take place, tilted positioning of the clamp on DNA restricting movement more than expected, or rotational movement coupled with translation, that latter of which was observed for PCNA [69]. Individually, these conditions do not create enough drag to account for the low diffusion rate calculated here, however, perhaps a combination of these scenarios, or the formation of secondary structure in the ssDNA region, could explain the friction that WT β clamp appears to experience in our experiments. Also, we observe clamps sliding on the single-stranded regions of the DNA about half of the time.

Crystallographic studies suggest that the space between the clamp inner pore residues and the incoming DNA is occupied by two or three water layers, facilitating the clamp to slide or “water skate” along dsDNA [23, 30, 43]. In this hypothesis, the energetic expense from breaking and reforming electrostatic interactions is reduced while still enabling β to glide along dsDNA rapidly [43].

Our results show that β R24A and β R24E variants drift on DNA more than 6.5-fold faster than WT β. The increase in diffusion rate is expected for R24E due to repulsions between the glutamate residue and the negatively charged DNA phosphate backbone. This observation for β R24A suggests that the electrostatic interactions of the positively charged residues slow sliding. Moreover, both R24 clamp variants have significantly shorter lifetimes on DNA relative to WT β. These results are consistent with DNA loading assays in the absence of polymerase, with β R24A/E variants dissociating from DNA faster than WT β. In the optical tweezers, the continuous flow could cause proteins to dissociate faster in real time, especially those bound more weakly. We also observed the γ complex binding to ssDNA regions (Figure 8); other studies found that the γ complex had a significantly higher affinity for a 50-mer ssDNA relative to p/t DNA that contained a ssDNA region with the same 50-mer sequence [39].

It may be that removal of the positive charge is the major contributor to the effect on diffusion observed with β R24 variants, rather than the repulsion introduced by the mutation R24E. Mutations to R24 led to a loss in attractive ionic interactions that are not compensated by the DNA contacts with other polar and positively charge residues still present inside the clamp pore, reinforcing the importance of the R24 residue to the β:DNA interaction as it pertains to loading and diffusion. A number of positively charged residues located at precise positions in the lining of the β clamp central pore are necessary for attractive, ionic interactions with the DNA phosphate backbone to stabilize DNA for efficient loading of the clamp and to modulate clamp movement along DNA.

### Accession IDs

- â clamp: UniProt P0A988
- ã truncation of DnaX: UniProt P06710
- ä: UniProt P28630
- ä′: UniProt P28631
- SSB: UniProt P0AGE0

## ACKNOWLEDGMENTS

This work was supported by the National Institutes of Health (R01GM123239 to P.J.B. and D.M.K.) and the National Science Foundation (MCB-1615946 to P.J.B). M.L.L. was supported by a GEM Fellowship and an Albert Sebag PhD ’02 Graduate Fellowship.

