## Supplementary Information for "Mutation of charged inner pore residues reduce *E. coli* β clamp residency and increase sliding rates on DNA"

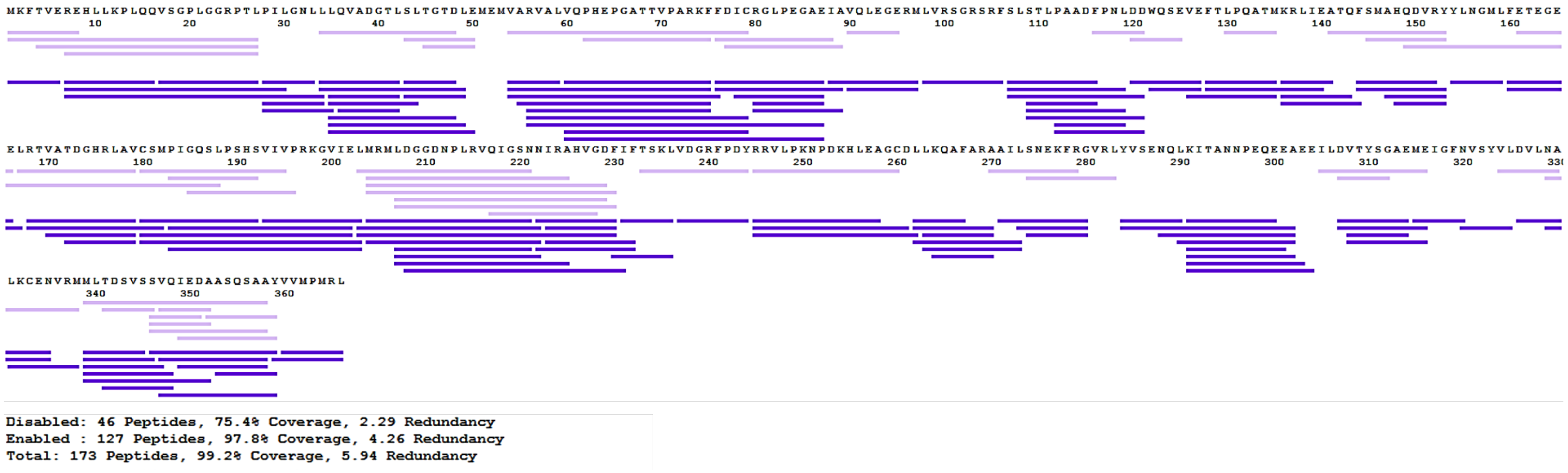

Figure S1. Peptide coverage map for WT β.

### Table S1. Relative Percent Deuterium Incorporation

| | | | | WT β clamp | | | | | | | β Q149A | | | | | |
|---|---|---|---|---|---|---|---|---|---|---|---|---|---|---|---|---|
| Sequence | Start | End | | 10 s | 1 min | 10 min | 1 h | 4 h | 6 h | | 10 s | 1 min | 10 min | 1 h | 4 h | 6 h |
| MKFTVE | 1 | 6 | | 0.0639 | 0.1289 | 0.2207 | 0.3061 | 0.3028 | 0.3142 | | 0.0479 | 0.1258 | 0.2102 | 0.2952 | 0.2953 | 0.3050 |
| MKFTVEREHLLKPLQQVSGPLGGRPTL | 1 | 27 | | 0.2554 | 0.3254 | 0.4099 | 0.4867 | 0.5314 | 0.5413 | | 0.2517 | 0.3202 | 0.4031 | 0.4759 | 0.5144 | 0.5402 |
| TVEREHLLKPLQQVSGPLGGRPTL | 4 | 27 | | 0.2980 | 0.3749 | 0.4578 | 0.5312 | 0.5812 | 0.5915 | | 0.2923 | 0.3653 | 0.4432 | 0.5165 | 0.5601 | 0.5878 |
| REHLLKPLQQ | 7 | 16 | | 0.0721 | 0.1391 | 0.2450 | 0.3299 | 0.3936 | 0.4163 | | 0.0571 | 0.1323 | 0.2230 | 0.3162 | 0.3730 | 0.4015 |
| REHLLKPLQQVSGPLGGRPTL | 7 | 27 | | 0.3251 | 0.3925 | 0.4859 | 0.5690 | 0.6202 | 0.6318 | | 0.3166 | 0.3829 | 0.4721 | 0.5521 | 0.5989 | 0.6317 |
| REHLLKPLQQVSGPLGGRPTLPIL | 7 | 30 | | 0.3290 | 0.4005 | 0.5034 | 0.5959 | 0.6436 | 0.6553 | | 0.3191 | 0.3894 | 0.4909 | 0.5850 | 0.6204 | 0.6480 |
| REHLLKPLQQVSGPLGGRPTLPILGNLL | 7 | 34 | | 0.2700 | 0.3406 | 0.4631 | 0.5616 | 0.5967 | 0.6075 | | 0.2686 | 0.3379 | 0.4571 | 0.5460 | 0.5821 | 0.6062 |
| VSGPLGGRPTL | 17 | 27 | | 0.6060 | 0.6761 | 0.7173 | 0.7186 | 0.7172 | 0.7171 | | 0.5987 | 0.6527 | 0.6995 | 0.6983 | 0.6892 | 0.7128 |
| PILGNL | 28 | 33 | | 0.0381 | 0.1031 | 0.3929 | 0.5747 | 0.5919 | 0.5845 | | 0.0307 | 0.1000 | 0.3842 | 0.5602 | 0.5534 | 0.5684 |
| PILGNLL | 28 | 34 | | 0.0332 | 0.0750 | 0.3009 | 0.4486 | 0.4569 | 0.4527 | | 0.0229 | 0.0719 | 0.2930 | 0.4287 | 0.4287 | 0.4450 |
| PILGNLLL | 28 | 35 | | 0.0316 | 0.0735 | 0.2602 | 0.3969 | 0.4017 | 0.4066 | | 0.0242 | 0.0644 | 0.2632 | 0.3805 | 0.3843 | 0.3921 |
| LLQVADGTL | 34 | 42 | | 0.1609 | 0.1825 | 0.2940 | 0.3732 | 0.4032 | 0.4132 | | 0.1583 | 0.1786 | 0.2903 | 0.3619 | 0.3931 | 0.4148 |
| LQVADGTL | 35 | 42 | | 0.1775 | 0.2055 | 0.3276 | 0.4127 | 0.4456 | 0.4620 | | 0.1633 | 0.1899 | 0.3090 | 0.3917 | 0.4244 | 0.4538 |
| QVADGTL | 36 | 42 | | 0.2340 | 0.2641 | 0.4098 | 0.5032 | 0.5000 | 0.5045 | | 0.2142 | 0.2412 | 0.3843 | 0.4662 | 0.4708 | 0.4836 |
| LQVADGTLSL | 35 | 44 | | 0.1465 | 0.1725 | 0.2627 | 0.3341 | 0.3600 | 0.3803 | | 0.1453 | 0.1674 | 0.2640 | 0.3228 | 0.3527 | 0.3761 |
| LLQVADGTLSLTGTDL | 34 | 49 | | 0.1006 | 0.1338 | 0.2086 | 0.2820 | 0.2911 | 0.3240 | | 0.0913 | 0.1097 | 0.1938 | 0.2616 | 0.2870 | 0.3117 |
| LQVADGTLSLTGTD | 35 | 48 | | 0.1152 | 0.1267 | 0.1938 | 0.2446 | 0.2920 | 0.3120 | | 0.1110 | 0.1268 | 0.1918 | 0.2376 | 0.2755 | 0.3131 |
| LQVADGTLSLTGTDL | 35 | 49 | | 0.1070 | 0.1262 | 0.2240 | 0.2885 | 0.3191 | 0.3403 | | 0.1069 | 0.1289 | 0.2216 | 0.2868 | 0.3161 | 0.3412 |
| LQVADGTLSLTGTDLE | 35 | 50 | | 0.1084 | 0.1407 | 0.2610 | 0.3196 | 0.3511 | 0.3696 | | 0.1034 | 0.1405 | 0.2509 | 0.3145 | 0.3395 | 0.3699 |
| SLTGTD | 43 | 48 | | 0.0399 | 0.0386 | 0.0466 | 0.0617 | 0.1027 | 0.1232 | | 0.0304 | 0.0339 | 0.0390 | 0.0502 | 0.0890 | 0.1222 |
| SLTGTDL | 43 | 49 | | 0.0354 | 0.0476 | 0.1206 | 0.1748 | 0.2080 | 0.2237 | | 0.0273 | 0.0462 | 0.1169 | 0.1657 | 0.2011 | 0.2273 |
| VARVAL | 54 | 59 | | 0.1694 | 0.2189 | 0.2313 | 0.3374 | 0.4880 | 0.5162 | | 0.1541 | 0.2055 | 0.2349 | 0.3250 | 0.4722 | 0.5077 |
| VARVALVQPHEPGATTVPARKF | 54 | 75 | | 0.2628 | 0.3139 | 0.3403 | 0.3939 | 0.4807 | 0.4903 | | 0.2497 | 0.3054 | 0.3273 | 0.3806 | 0.4604 | 0.4894 |
| VARVALVQPHEPGATTVPARKFF | 54 | 76 | | 0.2409 | 0.2909 | 0.3124 | 0.3627 | 0.4414 | 0.4599 | | 0.2420 | 0.2917 | 0.3123 | 0.3633 | 0.4372 | 0.4646 |

| | | | | | | | | | | | | | | | |
|---|---|---|---|---|---|---|---|---|---|---|---|---|---|---|---|
| VARVALVQPHEPGATTVPARKFFDIC | 54 | 79 | | 0.1952 | 0.2380 | 0.2559 | 0.2984 | 0.3626 | 0.3747 | | 0.1954 | 0.2399 | 0.2583 | 0.2970 | 0.3556 | 0.3791 |
| VARVALVQPHEPGATTVPARKFFDICRGLPEGAE | 54 | 87 | | 0.2048 | 0.2582 | 0.2766 | 0.3115 | 0.3692 | 0.3814 | | 0.1982 | 0.2518 | 0.2691 | 0.3044 | 0.3539 | 0.3772 |
| ARVALVQPHEPGATTVPARKF | 55 | 75 | | 0.2774 | 0.3313 | 0.3617 | 0.4000 | 0.4723 | 0.4827 | | 0.2710 | 0.3231 | 0.3453 | 0.3923 | 0.4595 | 0.4912 |
| RVALVQPHEPGATTVPARKF | 56 | 75 | | 0.2843 | 0.3416 | 0.3654 | 0.4115 | 0.4883 | 0.4995 | | 0.2775 | 0.3367 | 0.3570 | 0.4029 | 0.4743 | 0.5012 |
| RVALVQPHEPGATTVPARKFFDIC | 56 | 79 | | 0.2199 | 0.2657 | 0.2818 | 0.3167 | 0.3703 | 0.3836 | | 0.2159 | 0.2625 | 0.2781 | 0.3084 | 0.3546 | 0.3794 |
| RVALVQPHEPGATTVPARKFFDICRGLPEGAE | 56 | 87 | | 0.2336 | 0.2801 | 0.3020 | 0.3304 | 0.3686 | 0.3933 | | 0.2183 | 0.2691 | 0.2919 | 0.3164 | 0.3595 | 0.3861 |
| VQPHEPGATTVPARKF | 60 | 75 | | 0.1967 | 0.2453 | 0.2691 | 0.3141 | 0.3767 | 0.3810 | | 0.1899 | 0.2398 | 0.2593 | 0.3020 | 0.3631 | 0.3809 |
| VQPHEPGATTVPARKFFDIC | 60 | 79 | | 0.1341 | 0.1679 | 0.1829 | 0.2103 | 0.2494 | 0.2574 | | 0.1265 | 0.1648 | 0.1780 | 0.2018 | 0.2410 | 0.2560 |
| VQPHEPGATTVPARKFFDICRGLPEGAE | 60 | 87 | | 0.1615 | 0.2167 | 0.2254 | 0.2533 | 0.2850 | 0.3014 | | 0.1565 | 0.2110 | 0.2240 | 0.2447 | 0.2764 | 0.3017 |
| FDICRGLPEGAE | 76 | 87 | | 0.1566 | 0.2093 | 0.2187 | 0.2349 | 0.2647 | 0.2766 | | 0.1529 | 0.2074 | 0.2161 | 0.2271 | 0.2512 | 0.2781 |
| FDICRGLPEGAEIA | 76 | 89 | | 0.1386 | 0.1851 | 0.1929 | 0.1998 | 0.2183 | 0.2264 | | 0.1347 | 0.1814 | 0.1889 | 0.1946 | 0.2032 | 0.2259 |
| ICRGLPEGAE | 78 | 87 | | 0.2129 | 0.2618 | 0.2925 | 0.3042 | 0.3573 | 0.3545 | | 0.2233 | 0.2887 | 0.3057 | 0.3111 | 0.3519 | 0.3557 |
| RGLPEGAE | 80 | 87 | | 0.2570 | 0.3464 | 0.3683 | 0.3899 | 0.4337 | 0.4375 | | 0.2532 | 0.3390 | 0.3574 | 0.3747 | 0.4094 | 0.4360 |
| RGLPEGAEIA | 80 | 89 | | 0.1992 | 0.2658 | 0.2799 | 0.2911 | 0.3173 | 0.3203 | | 0.1963 | 0.2636 | 0.2766 | 0.2876 | 0.2995 | 0.3222 |
| IAVQLEGERM | 88 | 97 | | 0.1758 | 0.2658 | 0.3815 | 0.4027 | 0.4177 | 0.4206 | | 0.1641 | 0.2525 | 0.3695 | 0.3885 | 0.3916 | 0.4177 |
| VQLEGERM | 90 | 97 | | 0.2520 | 0.3658 | 0.5161 | 0.5378 | 0.5596 | 0.5605 | | 0.2385 | 0.3500 | 0.4945 | 0.5232 | 0.5270 | 0.5517 |
| LVRSGRSRF | 98 | 106 | | 0.1733 | 0.1774 | 0.1781 | 0.1887 | 0.2047 | 0.2022 | | 0.1700 | 0.1688 | 0.1710 | 0.1792 | 0.1902 | 0.2050 |
| SLSTLPAADF | 107 | 116 | | 0.1386 | 0.2181 | 0.2771 | 0.3709 | 0.4752 | 0.4888 | | 0.1361 | 0.2115 | 0.2749 | 0.3676 | 0.4459 | 0.4779 |
| STLPAADF | 109 | 116 | | 0.1908 | 0.3088 | 0.3823 | 0.4608 | 0.4990 | 0.5086 | | 0.1889 | 0.3004 | 0.3732 | 0.4448 | 0.4769 | 0.5058 |
| SLSTLPAADFPNL | 107 | 119 | | 0.2400 | 0.3423 | 0.3931 | 0.4813 | 0.5658 | 0.5822 | | 0.2197 | 0.3254 | 0.3804 | 0.4640 | 0.5392 | 0.5722 |
| SLSTLPAADFPNLDD | 107 | 121 | | 0.2599 | 0.3459 | 0.3852 | 0.4578 | 0.5247 | 0.5413 | | 0.2452 | 0.3336 | 0.3766 | 0.4454 | 0.5093 | 0.5372 |
| STLPAADFPNL | 109 | 119 | | 0.2933 | 0.4125 | 0.4670 | 0.5180 | 0.5535 | 0.5631 | | 0.2773 | 0.4096 | 0.4596 | 0.5091 | 0.5253 | 0.5551 |
| STLPAADFPNLDD | 109 | 121 | | 0.3139 | 0.4170 | 0.4467 | 0.4924 | 0.5138 | 0.5200 | | 0.2997 | 0.3967 | 0.4403 | 0.4796 | 0.4955 | 0.5190 |
| PAADFPNL | 112 | 119 | | 0.4339 | 0.5618 | 0.6185 | 0.7075 | 0.7481 | 0.7493 | | 0.4075 | 0.5453 | 0.6084 | 0.6803 | 0.7200 | 0.7376 |
| PAADFPNLDD | 112 | 121 | | 0.4148 | 0.4982 | 0.5359 | 0.5842 | 0.6222 | 0.6153 | | 0.3878 | 0.4752 | 0.5088 | 0.5622 | 0.5822 | 0.6017 |
| DDWQSEVE | 120 | 127 | | 0.3709 | 0.5237 | 0.6157 | 0.6299 | 0.6299 | 0.6313 | | 0.3676 | 0.5146 | 0.5921 | 0.6176 | 0.6159 | 0.6204 |
| WQSEVE | 122 | 127 | | 0.2924 | 0.4299 | 0.5569 | 0.5868 | 0.5891 | 0.5814 | | 0.2778 | 0.4063 | 0.5363 | 0.5604 | 0.5583 | 0.5735 |
| VEFTLPQATM | 126 | 135 | | 0.1752 | 0.3067 | 0.3769 | 0.3822 | 0.3877 | 0.3837 | | 0.1655 | 0.2914 | 0.3686 | 0.3682 | 0.3679 | 0.3768 |

| | | | | | | | | | | | | | | | |
|---|---|---|---|---|---|---|---|---|---|---|---|---|---|---|---|
| FTLPQATM | 128 | 135 | | 0.2014 | 0.3295 | 0.3613 | 0.3652 | 0.3666 | 0.3666 | | 0.1974 | 0.3141 | 0.3533 | 0.3520 | 0.3543 | 0.3691 |
| FTLPQATMKRLIE | 128 | 140 | | 0.1094 | 0.1910 | 0.2031 | 0.2016 | 0.2017 | 0.2043 | | 0.1027 | 0.1751 | 0.1935 | 0.1929 | 0.1941 | 0.2042 |
| KRLIEA | 136 | 141 | | 0.0305 | 0.0327 | 0.0382 | 0.0387 | 0.0389 | 0.0621 | | 0.0123 | 0.0049 | 0.0105 | 0.0162 | 0.0055 | 0.0260 |
| KRLIEATQ | 136 | 143 | | 0.0288 | 0.0260 | 0.0306 | 0.0340 | 0.0498 | 0.0630 | | 0.0253 | 0.0240 | 0.0281 | 0.0292 | 0.0350 | 0.0442 |
| KRLIEATQF | 136 | 144 | | 0.0670 | 0.1153 | 0.1193 | 0.1241 | 0.1434 | 0.1493 | | 0.0937 | 0.1092 | 0.1099 | 0.1125 | 0.1151 | 0.1260 |
| FSMAHQDVR | 144 | 152 | | 0.4173 | 0.4520 | 0.4953 | 0.4992 | 0.4976 | 0.4869 | | 0.4268 | 0.4312 | 0.4586 | 0.4971 | 0.5059 | 0.5029 |
| FSMAHQDVRY | 144 | 153 | | 0.3971 | 0.4603 | 0.5113 | 0.5155 | 0.5138 | 0.5023 | | 0.3845 | 0.3947 | 0.4485 | 0.5063 | 0.5146 | 0.5152 |
| AHQDVRY | 147 | 153 | | 0.5277 | 0.5581 | 0.5981 | 0.6107 | 0.6123 | 0.5957 | | 0.5421 | 0.5585 | 0.5653 | 0.5842 | 0.6399 | 0.5912 |
| HQDVRY | 148 | 153 | | 0.5060 | 0.5258 | 0.5126 | 0.5268 | 0.5265 | 0.5053 | | 0.4836 | 0.4877 | 0.4837 | 0.4858 | 0.4760 | 0.4954 |
| YLNGML | 154 | 159 | | 0.1311 | 0.1422 | 0.1906 | 0.2582 | 0.2695 | 0.2648 | | 0.0695 | 0.1106 | 0.1288 | 0.1812 | 0.2290 | 0.2382 |
| FETEGEE | 160 | 166 | | 0.1919 | 0.2410 | 0.3509 | 0.4079 | 0.4132 | 0.4119 | | 0.1610 | 0.2170 | 0.2873 | 0.3819 | 0.3918 | 0.4036 |
| FETEGEEL | 160 | 167 | | 0.1270 | 0.1532 | 0.2431 | 0.2935 | 0.2970 | 0.2952 | | 0.1136 | 0.1332 | 0.1930 | 0.2755 | 0.2771 | 0.2906 |
| LRTVATDGHRLAV | 167 | 179 | | 0.0625 | 0.0707 | 0.0895 | 0.1022 | 0.1149 | 0.1144 | | 0.0543 | 0.0561 | 0.0715 | 0.0863 | 0.0902 | 0.0960 |
| RTVATDGHRLAV | 168 | 179 | | 0.0629 | 0.0769 | 0.0941 | 0.1068 | 0.1199 | 0.1164 | | 0.0417 | 0.0548 | 0.0719 | 0.0842 | 0.0880 | 0.0913 |
| RTVATDGHRLAVCSM | 168 | 182 | | 0.0471 | 0.0587 | 0.0867 | 0.1190 | 0.1318 | 0.1331 | | 0.0327 | 0.0454 | 0.0650 | 0.0954 | 0.1123 | 0.1173 |
| VATDGHRLAV | 170 | 179 | | 0.0709 | 0.0915 | 0.1203 | 0.1364 | 0.1531 | 0.1557 | | 0.0546 | 0.0701 | 0.0947 | 0.1113 | 0.1139 | 0.1297 |
| TDGHRLAV | 172 | 179 | | 0.0852 | 0.1104 | 0.1333 | 0.1592 | 0.1787 | 0.1832 | | 0.0458 | 0.0689 | 0.0962 | 0.1199 | 0.1269 | 0.1355 |
| CSMPIGQSLPSHS | 180 | 192 | | 0.3230 | 0.3586 | 0.4121 | 0.4735 | 0.5132 | 0.5211 | | 0.3134 | 0.3489 | 0.3883 | 0.4491 | 0.4823 | 0.4959 |
| CSMPIGQSLPSHSVIVPRKGVIE | 180 | 202 | | 0.1973 | 0.2409 | 0.2851 | 0.3237 | 0.3637 | 0.3728 | | 0.1907 | 0.2358 | 0.2707 | 0.3026 | 0.3346 | 0.3580 |
| CSMPIGQSLPSHSVIVPRKGVIE L | 180 | 203 | | 0.1812 | 0.2215 | 0.2641 | 0.3023 | 0.3371 | 0.3473 | | 0.1741 | 0.2145 | 0.2470 | 0.2797 | 0.3121 | 0.3303 |
| PIGQSLPSHSVIVPRKGVIE | 183 | 202 | | 0.2275 | 0.2805 | 0.3306 | 0.3675 | 0.3885 | 0.3927 | | 0.2216 | 0.2746 | 0.3127 | 0.3479 | 0.3719 | 0.3883 |
| PIGQSLPSHSVIVPRKGVIEL | 183 | 203 | | 0.2064 | 0.2542 | 0.3017 | 0.3379 | 0.3588 | 0.3610 | | 0.1982 | 0.2464 | 0.2807 | 0.3153 | 0.3406 | 0.3542 |
| VIVPRKGVIEL | 193 | 203 | | 0.0895 | 0.1489 | 0.1892 | 0.2079 | 0.2416 | 0.2518 | | 0.1012 | 0.1554 | 0.1894 | 0.1970 | 0.2267 | 0.2409 |
| LMRMLDGGDNPLRVQIGSNN | 203 | 222 | | 0.2209 | 0.2603 | 0.2966 | 0.3569 | 0.4320 | 0.4603 | | 0.2168 | 0.2497 | 0.2904 | 0.3432 | 0.4219 | 0.4684 |
| MRMLDGGDNPLRVQIGSN | 204 | 221 | | 0.2291 | 0.2494 | 0.2827 | 0.3474 | 0.4332 | 0.4614 | | 0.2248 | 0.2419 | 0.2772 | 0.3340 | 0.4233 | 0.4610 |
| MRMLDGGDNPLRVQIGSNN | 204 | 222 | | 0.2248 | 0.2645 | 0.2993 | 0.3566 | 0.4395 | 0.4621 | | 0.2195 | 0.2565 | 0.2925 | 0.3440 | 0.4264 | 0.4639 |
| MRMLDGGDNPLRVQIGSNNIRAH VGD | 204 | 229 | | 0.2016 | 0.2308 | 0.2559 | 0.3062 | 0.3796 | 0.4038 | | 0.2016 | 0.2263 | 0.2502 | 0.3001 | 0.3716 | 0.4062 |
| MRMLDGGDNPLRVQIGSNNIRAH VGDF | 204 | 230 | | 0.1754 | 0.1984 | 0.2264 | 0.2806 | 0.3640 | 0.3879 | | 0.1724 | 0.1929 | 0.2228 | 0.2757 | 0.3508 | 0.3883 |

| | | | | | | | | | | | | | | | |
|---|---|---|---|---|---|---|---|---|---|---|---|---|---|---|---|
| LDGGDNPLRVQIGSN | 207 | 221 | | 0.2516 | 0.2583 | 0.2652 | 0.3121 | 0.3989 | 0.4229 | | 0.2466 | 0.2469 | 0.2547 | 0.2968 | 0.3812 | 0.4111 |
| LDGGDNPLRVQIGSNN | 207 | 222 | | 0.2449 | 0.2771 | 0.2851 | 0.3288 | 0.4082 | 0.4313 | | 0.2402 | 0.2645 | 0.2742 | 0.3136 | 0.3865 | 0.4243 |
| LDGGDNPLRVQIGSNNIRA | 207 | 225 | | 0.2180 | 0.2434 | 0.2488 | 0.2857 | 0.3553 | 0.3749 | | 0.2019 | 0.2229 | 0.2322 | 0.2628 | 0.3307 | 0.3584 |
| LMRMLDGGDNPLRVQIGSNNIRA HVGDF | 203 | 230 | | 0.1733 | 0.1966 | 0.2234 | 0.2805 | 0.3721 | 0.3943 | | 0.1706 | 0.1889 | 0.2184 | 0.2732 | 0.3551 | 0.3945 |
| DGGDNPLRVQIGSNNIRAHVGDF I | 208 | 231 | | 0.1852 | 0.2046 | 0.2166 | 0.2580 | 0.3394 | 0.3665 | | 0.1781 | 0.1911 | 0.2029 | 0.2457 | 0.3179 | 0.3576 |
| NIRAHVGDF | 222 | 230 | | 0.1547 | 0.1532 | 0.1625 | 0.2137 | 0.2987 | 0.3117 | | 0.1561 | 0.1477 | 0.1625 | 0.2112 | 0.2856 | 0.3071 |
| NIRAHVGDFIF | 222 | 232 | | 0.0931 | 0.0999 | 0.1110 | 0.1526 | 0.2226 | 0.2320 | | 0.0997 | 0.1005 | 0.1089 | 0.1525 | 0.2091 | 0.2347 |
| IRAHVGDF | 223 | 230 | | 0.1480 | 0.1541 | 0.1656 | 0.2227 | 0.3096 | 0.3298 | | 0.1485 | 0.1526 | 0.1614 | 0.2205 | 0.2973 | 0.3312 |
| IRAHVGDFIF | 223 | 232 | | 0.0978 | 0.0980 | 0.1075 | 0.1527 | 0.2243 | 0.2374 | | 0.0926 | 0.0927 | 0.1097 | 0.1513 | 0.2198 | 0.2407 |
| FIFTSKL | 230 | 236 | | 0.0241 | 0.0329 | 0.0581 | 0.1281 | 0.1682 | 0.1885 | | 0.0157 | 0.0130 | 0.0546 | 0.1315 | 0.1640 | 0.1801 |
| IFTSKL | 231 | 236 | | 0.0261 | 0.0260 | 0.0568 | 0.1370 | 0.1920 | 0.2025 | | 0.0178 | 0.0258 | 0.0597 | 0.1395 | 0.1889 | 0.2097 |
| VDGRFPDY | 237 | 244 | | 0.3662 | 0.4819 | 0.5158 | 0.5151 | 0.5162 | 0.5057 | | 0.2753 | 0.4174 | 0.5030 | 0.5075 | 0.5033 | 0.5076 |
| RRVLPKNPDKHLEA | 245 | 258 | | 0.2138 | 0.2786 | 0.3900 | 0.4339 | 0.4663 | 0.4619 | | 0.2033 | 0.2453 | 0.3411 | 0.4034 | 0.4387 | 0.4534 |
| RRVLPKNPDKHLEAGCD | 245 | 261 | | 0.2275 | 0.2985 | 0.3870 | 0.4268 | 0.4566 | 0.4611 | | 0.2016 | 0.2546 | 0.3328 | 0.3827 | 0.4191 | 0.4391 |
| RRVLPKNPDKHLEAGCDL | 245 | 262 | | 0.2024 | 0.2961 | 0.3906 | 0.4285 | 0.4567 | 0.4613 | | 0.1891 | 0.2697 | 0.3513 | 0.3986 | 0.4292 | 0.4517 |
| LLKQAF | 262 | 267 | | 0.0155 | 0.0194 | 0.0294 | 0.0841 | 0.1365 | 0.1458 | | 0.0097 | 0.0111 | 0.0263 | 0.0695 | 0.1237 | 0.1507 |
| LLKQAFARA | 262 | 270 | | 0.0300 | 0.0346 | 0.0450 | 0.0838 | 0.1255 | 0.1347 | | 0.0249 | 0.0316 | 0.0461 | 0.0772 | 0.1179 | 0.1295 |
| LLKQAFARAAIL | 262 | 273 | | 0.0097 | 0.0159 | 0.0242 | 0.0575 | 0.1015 | 0.1274 | | 0.0100 | 0.0183 | 0.0320 | 0.0646 | 0.1111 | 0.1355 |
| LKQAFARA | 263 | 270 | | 0.0326 | 0.0402 | 0.0539 | 0.0875 | 0.1337 | 0.1333 | | 0.0304 | 0.0422 | 0.0577 | 0.0859 | 0.1216 | 0.1394 |
| LKQAFARAAIL | 263 | 273 | | 0.0207 | 0.0251 | 0.0345 | 0.0596 | 0.1058 | 0.1226 | | 0.0172 | 0.0248 | 0.0258 | 0.0537 | 0.1007 | 0.1189 |
| KQAFARA | 264 | 270 | | 0.0433 | 0.0491 | 0.0636 | 0.0935 | 0.1371 | 0.1507 | | 0.0290 | 0.0442 | 0.0586 | 0.0928 | 0.1196 | 0.1395 |
| AILSNEKFRG | 271 | 280 | | 0.3835 | 0.4381 | 0.5091 | 0.5468 | 0.5649 | 0.5605 | | 0.3731 | 0.4182 | 0.4912 | 0.5290 | 0.5512 | 0.5654 |
| LSNEKFRG | 273 | 280 | | 0.5072 | 0.5640 | 0.6544 | 0.6958 | 0.7127 | 0.7088 | | 0.4939 | 0.5405 | 0.6301 | 0.6630 | 0.6647 | 0.6954 |
| SNEKFRG | 274 | 280 | | 0.5120 | 0.5930 | 0.6300 | 0.6299 | 0.6401 | 0.6278 | | 0.4908 | 0.5469 | 0.6011 | 0.6054 | 0.6018 | 0.6097 |
| SNEKFRGVRL | 274 | 283 | | 0.2720 | 0.3153 | 0.3456 | 0.3533 | 0.3631 | 0.3507 | | 0.2865 | 0.3265 | 0.3589 | 0.3626 | 0.3713 | 0.3787 |
| YVSENQL | 284 | 290 | | 0.1365 | 0.2142 | 0.2667 | 0.3419 | 0.4016 | 0.4104 | | 0.1262 | 0.2082 | 0.2602 | 0.3421 | 0.3928 | 0.4169 |
| YVSENQLKITANNPEQEEA | 284 | 302 | | 0.0975 | 0.1848 | 0.2723 | 0.3279 | 0.3760 | 0.3917 | | 0.0903 | 0.1795 | 0.2669 | 0.3264 | 0.3692 | 0.3969 |
| NQLKITANNPEQEEA | 288 | 302 | | 0.0749 | 0.1514 | 0.2509 | 0.2971 | 0.3269 | 0.3401 | | 0.0667 | 0.1521 | 0.2457 | 0.2858 | 0.3203 | 0.3439 |

| | | | | | | | | | | | | | | | |
|---|---|---|---|---|---|---|---|---|---|---|---|---|---|---|---|
| KITANNPEQE | 291 | 300 | | 0.0915 | 0.2258 | 0.3867 | 0.4540 | 0.4740 | 0.4696 | | 0.0876 | 0.2288 | 0.3884 | 0.4411 | 0.4621 | 0.4705 |
| LKITANNPEQEEA | 290 | 302 | | 0.0872 | 0.1812 | 0.2960 | 0.3451 | 0.3870 | 0.3966 | | 0.0795 | 0.1734 | 0.2885 | 0.3358 | 0.3822 | 0.3958 |
| KITANNPEQEE | 291 | 301 | | 0.0956 | 0.2091 | 0.3584 | 0.4184 | 0.4509 | 0.4621 | | 0.0872 | 0.2117 | 0.3569 | 0.4052 | 0.4462 | 0.4650 |
| KITANNPEQEEA | 291 | 302 | | 0.0914 | 0.1977 | 0.3217 | 0.3805 | 0.4233 | 0.4371 | | 0.0771 | 0.1882 | 0.3106 | 0.3655 | 0.4059 | 0.4325 |
| KITANNPEQEEAE | 291 | 303 | | 0.0823 | 0.1773 | 0.2929 | 0.3461 | 0.3847 | 0.3972 | | 0.0773 | 0.1739 | 0.2883 | 0.3345 | 0.3760 | 0.4005 |
| KITANNPEQEEAEE | 291 | 304 | | 0.0782 | 0.1637 | 0.2729 | 0.3185 | 0.3599 | 0.3652 | | 0.0704 | 0.1652 | 0.2662 | 0.3149 | 0.3469 | 0.3706 |
| DVTYSGAE | 307 | 314 | | 0.3409 | 0.5348 | 0.6178 | 0.6208 | 0.6218 | 0.6098 | | 0.3310 | 0.5105 | 0.6049 | 0.6055 | 0.5983 | 0.6136 |
| VTYSGAE | 308 | 314 | | 0.4255 | 0.5896 | 0.6312 | 0.6343 | 0.6428 | 0.6405 | | 0.3882 | 0.5656 | 0.6083 | 0.6152 | 0.6106 | 0.6117 |
| DVTYSGAEME | 307 | 316 | | 0.2890 | 0.4858 | 0.5570 | 0.5666 | 0.5744 | 0.5729 | | 0.2812 | 0.4647 | 0.5510 | 0.5564 | 0.5600 | 0.5786 |
| VTYSGAEME | 308 | 316 | | 0.3243 | 0.5023 | 0.5558 | 0.5547 | 0.5703 | 0.5698 | | 0.3134 | 0.4819 | 0.5418 | 0.5446 | 0.5496 | 0.5741 |
| MEIGFN | 315 | 320 | | 0.0361 | 0.1467 | 0.2197 | 0.3039 | 0.3173 | 0.3090 | | 0.0226 | 0.0957 | 0.2115 | 0.2909 | 0.2942 | 0.3213 |
| NVSYVL | 320 | 325 | | 0.0889 | 0.2004 | 0.3133 | 0.3342 | 0.3718 | 0.3904 | | 0.0773 | 0.1694 | 0.2758 | 0.3162 | 0.3440 | 0.3697 |
| DVLNALKCEN | 326 | 335 | | 0.2522 | 0.3152 | 0.3434 | 0.3549 | 0.4131 | 0.4719 | | 0.2604 | 0.3071 | 0.3220 | 0.3532 | 0.3940 | 0.4448 |
| LKCENVRM | 331 | 338 | | 0.2036 | 0.2446 | 0.2516 | 0.2656 | 0.2534 | 0.2555 | | 0.1816 | 0.2144 | 0.2414 | 0.2434 | 0.2407 | 0.2453 |
| MLTDSVS | 339 | 345 | | 0.1671 | 0.2943 | 0.4936 | 0.5621 | 0.6039 | 0.6016 | | 0.1471 | 0.2425 | 0.4614 | 0.5289 | 0.5812 | 0.5918 |
| MLTDSVSS | 339 | 346 | | 0.2429 | 0.3516 | 0.5133 | 0.5750 | 0.6104 | 0.6095 | | 0.2271 | 0.3102 | 0.4890 | 0.5489 | 0.5932 | 0.6064 |
| MLTDSVSSV | 339 | 347 | | 0.2208 | 0.3165 | 0.4586 | 0.5114 | 0.5484 | 0.5385 | | 0.2065 | 0.2756 | 0.4410 | 0.4799 | 0.5124 | 0.5270 |
| MLTDSVSSVQ | 339 | 348 | | 0.1754 | 0.2554 | 0.3807 | 0.4304 | 0.4571 | 0.4548 | | 0.1649 | 0.2293 | 0.3676 | 0.4087 | 0.4442 | 0.4546 |
| MLTDSVSSVQIEDA | 339 | 352 | | 0.1358 | 0.1940 | 0.2728 | 0.3304 | 0.3787 | 0.3821 | | 0.1302 | 0.1753 | 0.2734 | 0.3037 | 0.3550 | 0.3790 |
| TDSVSSVQ | 341 | 348 | | 0.2077 | 0.2834 | 0.3939 | 0.4177 | 0.4063 | 0.4013 | | 0.1892 | 0.2516 | 0.3878 | 0.4040 | 0.3964 | 0.4132 |
| SVQIEDAASQSAAY | 346 | 359 | | 0.1809 | 0.2577 | 0.3078 | 0.3258 | 0.3581 | 0.3792 | | 0.1641 | 0.2442 | 0.2884 | 0.3085 | 0.3308 | 0.3692 |
| VQIEDAASQSAA | 347 | 358 | | 0.2081 | 0.2930 | 0.3451 | 0.3801 | 0.4207 | 0.4232 | | 0.1954 | 0.2807 | 0.3306 | 0.3649 | 0.3819 | 0.4054 |
| VQIEDAASQSAAY | 347 | 359 | | 0.1929 | 0.2679 | 0.3129 | 0.3480 | 0.3830 | 0.3941 | | 0.1812 | 0.2566 | 0.3045 | 0.3333 | 0.3538 | 0.3747 |
| IEDAASQSAA | 349 | 358 | | 0.2330 | 0.3234 | 0.3846 | 0.4211 | 0.4577 | 0.4757 | | 0.2123 | 0.3041 | 0.3709 | 0.3980 | 0.4255 | 0.4562 |
| ASQSAAY | 353 | 359 | | 0.2693 | 0.3568 | 0.3703 | 0.3783 | 0.3687 | 0.3725 | | 0.2600 | 0.3442 | 0.3599 | 0.3624 | 0.3622 | 0.3642 |
| YVVMPMRL | 359 | 366 | | 0.2283 | 0.3882 | 0.5479 | 0.5635 | 0.5632 | 0.5662 | | 0.2134 | 0.3233 | 0.4653 | 0.5472 | 0.5539 | 0.5665 |
| VVMPMRL | 360 | 366 | | 0.3049 | 0.5045 | 0.6871 | 0.7088 | 0.7227 | 0.7118 | | 0.2683 | 0.4086 | 0.5845 | 0.6742 | 0.6916 | 0.6960 |
